# Towards selective-alignment: Bridging the accuracy gap between alignment-based and alignment-free transcript quantification

**DOI:** 10.1101/138800

**Authors:** Hirak Sarkar, Mohsen Zakeri, Laraib Malik, Rob Patro

## Abstract

**Motivation:** We introduce an algorithm for selectively aligning high-throughput sequencing reads to a transcriptome, with the goal of improving transcript-level quantification. This algorithm attempts to bridge the gap between fast “mapping” algorithms and more traditional alignment procedures.

**Results:** We adopt a hybrid approach that is able to increase mapping accuracy while still retaining much of the efficiency of fast mapping algorithms. To achieve this, we introduce a new approach that explores the candidate search space with high sensitivity as well as a collection of carefully-engineered heuristics to efficiently filter these candidates. Additionally, unlike the strategies adopted in most aligners which first align the ends of paired-end reads independently, we introduce a notion of co-mapping. This procedure exploits relevant information between the “hits” from the left and right ends of paired-end reads before full alignments or mappings for each are generated, which improves the efficiency of filtering likely-spurious alignments. Finally, we demonstrate the utility of selective alignment in improving the accuracy of efficient transcript-level quantification from RNA-seq reads. Specifically, we show that selective-alignment is able to resolve certain complex mapping scenarios that can confound existing fast mapping procedures, while simultaneously eliminating spurious alignments that fast mapping approaches can produce.

**Availability:** Selective-alignment is implemented in C++11 as a part of *Salmon*, and is available as open source software, under GPL v3, at: https://github.com/COMBINE-lab/salmon/tree/selective-alignment

## 1. Introduction

Since the introduction of high-throughput, short read sequencing technologies, many algorithms and tools have been designed to tackle the problem of aligning short sequenced reads to a reference genome or transcriptome accurately and efficiently. While there exist “full-sensitivity” aligners (e.g. RazerS3 (Weese *et al.*, 2012), Masai (Siragusa *et al.*, 2013)) which guarantee to find all reference positions within a given edit-distance threshold of a read sequence, the most widely-used tools employ heuristic strategies to enable much faster alignment of reads in the typical case (i.e., only a small number of easy-to-find candidate locations exist for each alignment). The common procedure followed by these tools for aligning reads can be divided into two major steps. The first is finding potential alignment locations for the read using a pre-processed index that is generated from the reference genome or transcriptome. Then, in the second step, the potential locations are filtered, and reads are aligned to the positions that pass the initial filtering, based on a variety of heuristics. The exact method for generating the initial index varies for each tool. For example, tools like Bowtie (Langmead *et al.*, 2009), Bowtie2 ((Langmead and Salzberg, 2012)), BWA (Li and Durbin, 2009), and BWA-mem (Li, 2013) use Burrows-Wheeler transformation (BWT) based indices, whereas, k-mer based indices are used by tools such as Subread-aligner (Liao *et al.*, 2013), Maq (Li *et al.*, 2008), SNAP (Zaharia *et al.*, 2011), and GMAP and GSNAP (Wu and Nacu, 2010). Similarly, the heuristic for choosing the most probable locations is also different. However, each method is based on the principle of trying to find the reference loci that support the best (or near-best) alignment score between the read and the reference. Repeating this for a large number of reads comes with a considerable cost in terms of computation. Some tools, like STAR(Dobin *et al*., 2013), considerably speed up the alignment process by combining efficient heuristics with data structures (like the uncompressed suffix array) that trade working memory for exact pattern lookup speed. Recently, tools like HISAT (Kim *et al*., 2015) have also demonstrated that cache-friendly compressed indices (the hierarchical FM index in this case) can provide similarly efficient pattern search, even with a very moderate memory budget. The alignment of sequenced reads to the reference is the first step in pipelines leading to various downstream studies, such as estimation of transcript abundances and differential expression analysis, calculation of splicing rates (Shen *et al*., 2014; Vaquero-Garcia et al., 2016), and detection of fusion events (Nicorici et al., 2014; Davidson *et al*., 2015).

While alignment is a staple of many genomic analyses, it sometimes represents more information than is actually necessary to address the analysis at hand. For example, recent tools like *Sailfish* (Patro *et al*., 2014), *kallisto* (Bray *et al*., 2016), and Salmon (Patro *et al*., 2017), demonstrate that accurate quantification estimates can be obtained without all of the information encoded in traditional alignments. By avoiding traditional alignment procedure, these tools are much faster than their alignment-based counterparts. Furthermore, by building the mapping phase of the analysis directly into the quantification task, they dispense with the need to write, store, and read, large intermediate alignment files. However, these “mapping-based” tools, while highly-efficient, have the disadvantage of potentially losing sensitivity or specificity in certain cases where alignment-based methods would perform well. For example, in the presence of paralogous genes, with high sequence similarity, there is an increased probability that the mapping strategies employed by such tools, and the efficient heuristics upon which they rely, will mis-map reads between the paralogs (or return a more ambiguous set of mapping locations than an aligner, which expends effort to verify the returned alignments, would have) (Axtell, 2014). Similarly, in the case of *de novo* assemblies, poorly assembled contigs may have a larger number of mis-mapped reads due to lower sensitivity (here, the issue would be primarily due to aberrant exact matches masking the true origin of a read).

Other than suffering from spurious mappings, these fast mapping-based approaches can also miss true mappings of a read in rare cases where errors are positioned adversarially on the read. An obvious case of losing the true mapping is if a read contains no subsequence of sufficient length from the true transcript. In another case, the true mapping of the read might be lost from the set of potential mapping loci due to the greedy nature of the mapping procedures. For some reads, multiple positions might be found on the same transcript where the read maps. In such cases, improved heuristics are required to address these challenges.

In this paper, we present a novel algorithm, selective-alignment, that extends quasi-mapping to compute and store edit distance information where necessary. The reads for alignment are chosen based on certain criteria calculated during mapping. This strikes a balance between speed and accuracy; not compromising the superior speed of fast mapping algorithms, while also addressing some of the challenges mentioned above. Specifically, the motivation for selective-alignment is to enhance both the sensitivity and specificity of fast mapping algorithms by reducing or eliminating cases where spurious exact matches mask true mapping locations as well as cases where small exact matches support otherwise poor alignments. Selective-alignment algorithm is built atop the framework of *RapMap* (Srivastava *et al.*, 2016), which uses an index that combines a fixed-length prefix hash table and an uncompressed suffix array (Manber and Myers, 1993). We introduce a coverage-based consensus scheme to identify critical read candidates for which alignment is necessary. Further, we explored the challenging cases where the heuristics used by the fast mapping algorithms fail to locate the correct locations for a read, while the traditional aligners do not, and show that selective-alignment enables us to retain much of the improved accuracy of traditional aligners, but to do so more quickly. We also introduce filtering steps based on edit distance to further refine probable alignments in order to enhance quantification estimates (e.g., eliminating situations where the best mapping is still unlikely to represent the true origin of the read). In this work, we focus on the effect of selective-alignment in improving transcript quantification estimates, and we leave a thorough evaluation of the alignment qualities themselves as future work. In particular, evaluation of alignment qualities is considerably complicated by prevalent multi-mapping in the transcriptome.

## 2 Methods

The process of selective-alignment builds upon many of the basic data structures of Srivastava *et al.* (2016) yet there are a number of important algorithmic distinctions. Hence, we begin with a brief summary of the data structures backing the quasi-mapping implementation of *RapMap*. To start with, the index built on the transcriptome in selective-alignment is a combination of a suffix array and a hash table constructed from unique k-mers and suffix array intervals. The suffix array of a sequence, *T* — denoted *SA*(*T*)—is an array of starting positions of all suffixes from *T* in the original sequence. The values in the array are sorted lexicographically by the suffixes they represent. Therefore, all suffixes starting with the same prefix are located in adjacent positions of the suffix array. Formally, given a suffix array, *SA*(*T*) = Λ, constructed from the transcriptome sequence, *T*, we construct a hash table, *h*, that maps each k-mer, *κ*, to a suffix array interval, `I` (*κ*) = [*b*, *e*), if and only if all the suffixes within interval [*b*, *e*) contain the k-mer *κ* as a prefix. We define Λ[*i*] for every 0 ≤ *i* ≤ |Λ| to be the suffix *T* [*SA*[*i*]] (i.e., the suffix of *T* starting from position *SA*[*i*]). In selective-alignment’s index, in addition to suffix array intervals, we also store two extra pieces of information for each interval; the longest common prefix (LCP) and the k-safe-LCP corresponding to the interval. These are detailed below.

### 2.1 Defining and computing k-safe-LCPs

Here, we formally define the concept of k-safe-LCPs. The determination of k-safe-LCPs starts by labeling each suffix array interval with the length of its corresponding longest common prefix and the associated transcript set it represents. Formally, `LCP`(Λ[*b*], Λ[*e -* 1]) for an interval [*b, e*) is the length of the common prefix of the suffixes Λ[*b*] and Λ[*e -* 1].

Given k-mer *κ*, where *κ ∈****𝒦*** and ***𝒦*** is the set of all k-mers from the reference sequence *T*, and the related interval `I` (*κ*) = [*b, e*), for all *p ∈* [*b, e*), we consider each transcript *t* such that the suffix Λ[*p*] starts in transcript *t* in the concatenated text. Then, for this interval, we can construct a set *𝒞*^*κ*^ = {*t*_*i*_*, t*_*j*_ *, …* }, which denotes the set of distinct transcripts that appear in the suffix array interval, indicated by *κ*. We note that this notion discards duplicate appearances of the same transcript in this interval.

We now wish to define the notion of the k-safe-LCP of a suffix array interval. The k-safe-LCP of an interval `I` (*κ*) is the longest common prefix of the suffixes in the interval, where no k-mer occurring in this prefix belongs to a transcript not appearing in *𝒞*^*κ*^ (as defined above). We compute the k-safe-LCP for an interval indicated by k-mer *κ*_*i*_ iteratively. The initial length for the k-safe-LCP of the interval is *k*, length of a k-mer. We check, sequentially, each of the k-mers in the longest common prefix of the interval. For each new k-mer, the k-safe-LCP is increased by one character. We terminate the k-safe-LCP extension if any of the following conditions is encountered: (1) we reach the last k-mer contained in the LCP of this interval, (2) we encounter a k-mer *κ*_*j*_ such that *𝒞*^*κ*_*j*_^ ⊈ *𝒞* ^*κ*_*i*_^ or (3) we encounter a k-mer *κ*_*j*_ such that the reverse complement of *κ*_*j*_ appears elsewhere in transcriptome. When we encounter case (2) or (3), we call the k-mer *κ*_*j*_ an *intruder*. That is, the k-mer will potentially alter our belief about the set of potential transcripts to which a sequence containing this k-mer maps (by strictly expanding this set), or the orientation with which it maps to the transcriptome. We denote the k-safe-LCP of a particular interval `I` (*κ*_*i*_) as `k-safe-LCP`(`I` (*κ*_*i*_)). As shown in Figure 1, the k-safe-LCP determination for the top suffix array interval starts with matching k-mers within the longest common prefix. The k-mer “CAACG” maps to a suffix array interval labeled with (*t*_1_*, t*_2_). The next k-mer “AACGG”, on the other hand, maps to a suffix array interval (shaded in green) labeled with (*t*_1_*, t*_2_*, t*_3_), thereby implying the k-safe-LCP, shown as a dotted line. For each k-mer in the hash table, we store the length of the LCP and k-safe-LCP, along with the corresponding suffix array interval.

**Fig. 1:**
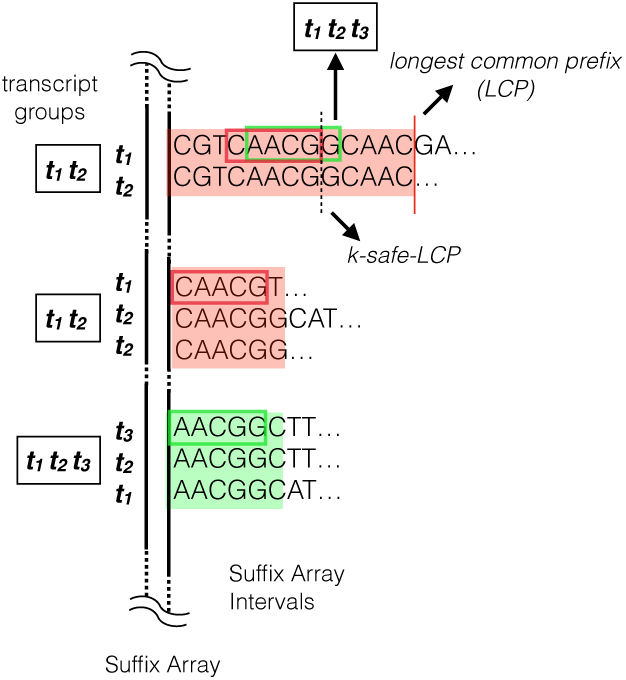
Calculation of k-safe-LCP from the suffix array data structure. The transcripts present in each suffix array interval determine the relevant transcript sets, and which k-mers will be considered as intruders. Detection of a k-mer that maps to suffix array interval labeled (*t*_1_*, t*_2_*, t*_3_) determines the k-safe-LCP here.

### 2.2 Discovering relevant suffix array intervals

As shown in Figure 2, the selective-alignment approach can be broken into three major steps: collecting suffix array intervals, co-mapping, and selecting the high quality mappings. Gathering the suffix array intervals for a query read closely follows the quasi-mapping approach. It involves iterating over the read from left to right and repeating two steps. First, hashing k-mer from the read sequence and then discovering the corresponding suffix array intervals. The process of k-mer lookup is aided by the k-safe-LCP stored in the index (discussed in Section 2.1). The inbuilt lexicographic ordering of the suffixes in the suffix array, and the computed k-safe-LCP values of intervals enable safely extending k-mers to longer matches without the possibility of masking potentially-informative substring matches. Given a matching k-mer, *κ*_*r*_, from the read sequence *r*, we extend the match to find the longest substring of the read that matches within `k-safe-LCP`(`I` (*κ*_*r*_)). The matched substring can be regarded as maximum mappable prefix (MMP) (Dobin *et al.*, 2013), that resides within the established k-safe-LCP. We call this a maximal mappable safe prefix (MMSP — eliding *k* where implied). For a k-mer, *κ*_*r*_, and interval, [*b, e*), we note that `k-safe-LCP`(`I` (*κ*_*r*_)) *≥ 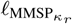*, where *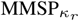* is the length of MMSP_*κ*_*r*, the MMSP between the read’s suffix starting with *κ*_*r*_ and the interval `I` (*κ*_*r*_). The next k-mer lookup starts from the 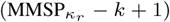-th position. By restricting our match extensions to reside within the MMSP, we ensure that we will not neglect to query any k-mer that might *expand* the set of potential transcripts where our read may map. We note here both the theoretical and practical relation between the MMSP matching procedure, and the concept of a uni-MEM, as introduced by Liu *et al.* (2016). The k-safe-LCP for suffix array intervals are closely related to the lengths of unipaths in the reference de Bruijn graph of order *k*. Thus, our procedure for finding MMSPs, that limits match extension by the k-safe-LCP, is similar to the uni-MEM seed generation procedure described in deBGA (Liu *et al.*, 2016), with the distinction that here, we only consider extending seeds in one direction.

**Fig. 2:**
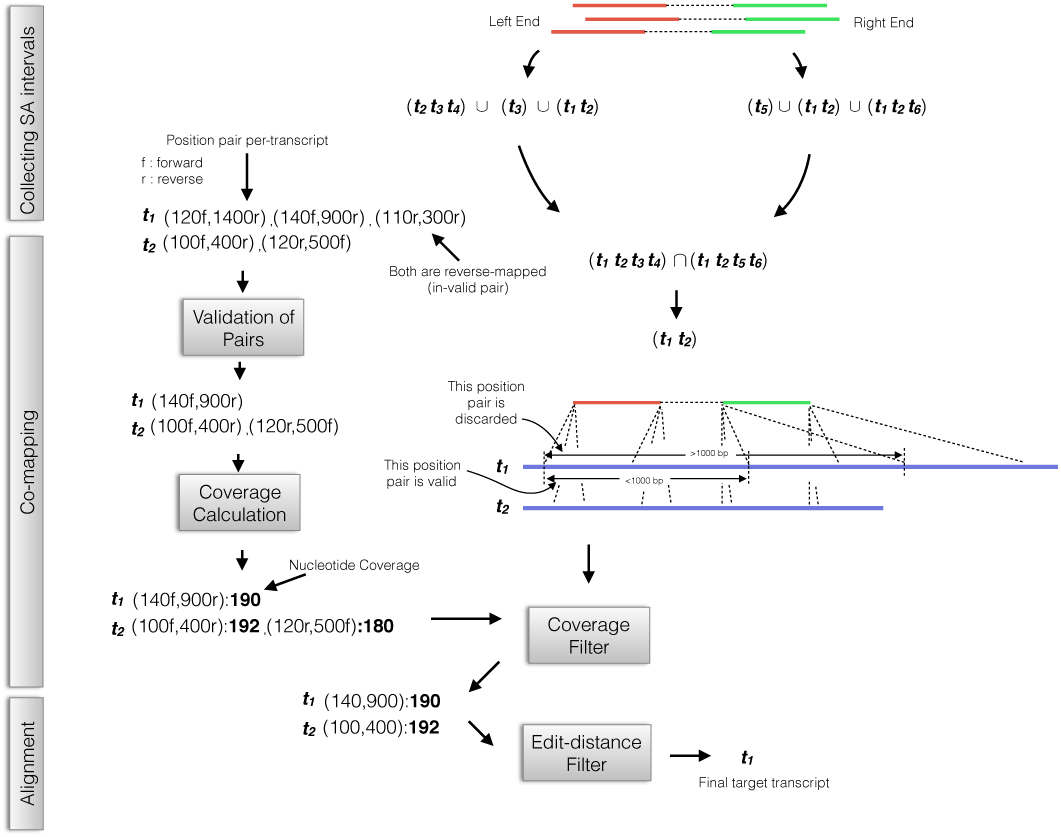
The three main steps of the selective-alignment process are demonstrated here. First, suffix array “hits” are collected. Then, in co-mapping, spurious mappings are removed by the orientation filter and then distance filter. At most a single locus per-transcript is selected based on the coverage filter. Finally, an edit-distance-based filter is used to select the valid target transcripts.

Given all the suffix array intervals collected for a read end (i.e. one end of a paired-end read), we take the *union* of all the transcripts they encode. Formally, if a read *r* maps to suffix array intervals labeled with *𝒞*^*r*_1_^ *, …, 𝒞*^*r*_*n*_^, then we consider all transcripts in the set *𝒞* ^*r*_1_^ *∪𝒞* ^*r*_2_^ *∪ … ∪𝒞* ^*r*_*n*_^, and the associated positions implied by the suffix array intervals. As shown in Figure 2; this step is done before co-mapping. We note that, in practice, we actually adopt a hybrid approach for collecting the suffix array intervals. Specifically, when the MMSP is only of length *k*, instead of moving to the next k-mer, we jump by |*r*|/10 nucleotides (where |*r*| is the read length) before looking up the next k-mer, otherwise (if *|*MMSP| > *k*), we skip by the MMSP length as described. This heuristic prevents us from performing excessive lookups in low-complexity and repetitive regions of the transcriptome. We observe that, in practice, the k-safe-LCP, and hence the MMSP lengths can be quite large (Figure 3).

**Fig. 3:**
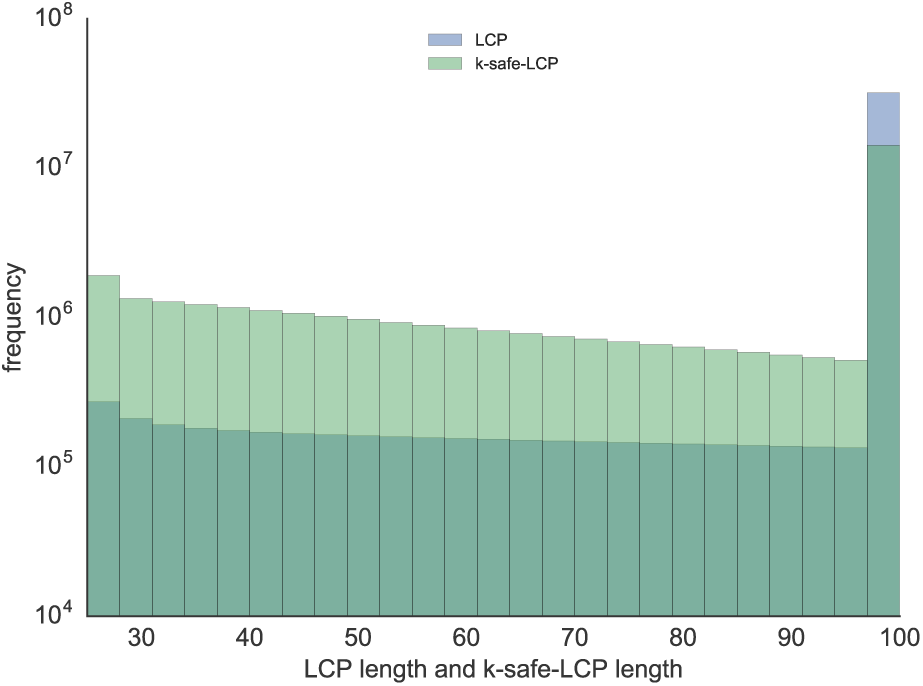
The distribution of k-safe-LCP lengths and LCP lengths are similar and tend to be large in practice (human transcriptome). Here, we truncate all lengths to a maximum value of 100 (so that any LCP or k-safe-LCP longer than 100 nucleotides is placed in the length 100 bin).

### 2.3 Co-Mapping

After collecting the suffix array intervals corresponding to left and right ends of the read, we wish to exploit the paired-end information in determining which potential mapping locations might be valid. Hence, from this step onward, we use the joint information for determining the position and target transcripts. Given the suffix array intervals for individual ends of a paired end read, the problem of aligning both ends of the pair poses a few challenges. First, a single read can map to multiple transcripts, and we wish to report all equally-best loci. Second, there can be multiple hits from a read on a single transcript (e.g., if a transcript contains repetitive sequence), and extra care must be taken to determine the correct mapping location. Finally, there may be hits that do not yield high-quality alignments (i.e. long exact matches that are nonetheless spurious). To address the first and third points, we employ an edit distance filter to discard spurious and sub-optimal alignments. To address the second challenge, we devise a consensus strategy to choose at most one unique position from each transcript.

Before applying the above mentioned strategy, we remove transcripts that do not contain hits from both the left and right ends of the read. Formally, given two ends of a read *r* as, *r*^*e*_1_^ and *r*^*e*_2_^, and the corresponding suffix array intervals labeled with *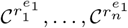* and *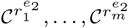* respectively, we only consider transcripts present in the set 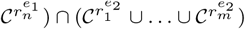 We further refine this set by checking the validity of the alignments these hits might support. Currently, we use two validity checks illustrated in Figure 2. First, we apply an orientation-based check, and second we employ a distance-based check. The orientation check removes potential mappings which have an orientation inconsistent with the underlying sequencing library type (e.g., both ends of a read mapping in the same orientation). The distance-based check removes potential alignments where the implied distance between the read ends is larger than a given, user-defined threshold (1, 000 nucleotides by default).

#### 2.3.1 Coverage based consensus

In selective-alignment, the potential positions on a transcript are scored by their individual coverage on the target transcript. Figure 4 depicts the mechanism of choosing the best postion on a transcript among multiple probable mapping to the same transcript. The coverage mechanism employed in selective-alignment makes use of the MMSP lengths collected during a prior step of the algorithm rather than simply counting k-mers. In Figure 4 the transcript *t*_2_ has two potential mapping position given the reads: 10 and 20, the coverage consensus mechanism selects 20 over position 10 due to higher coverage by tiling MMSPs on the read.

**Fig. 4:**
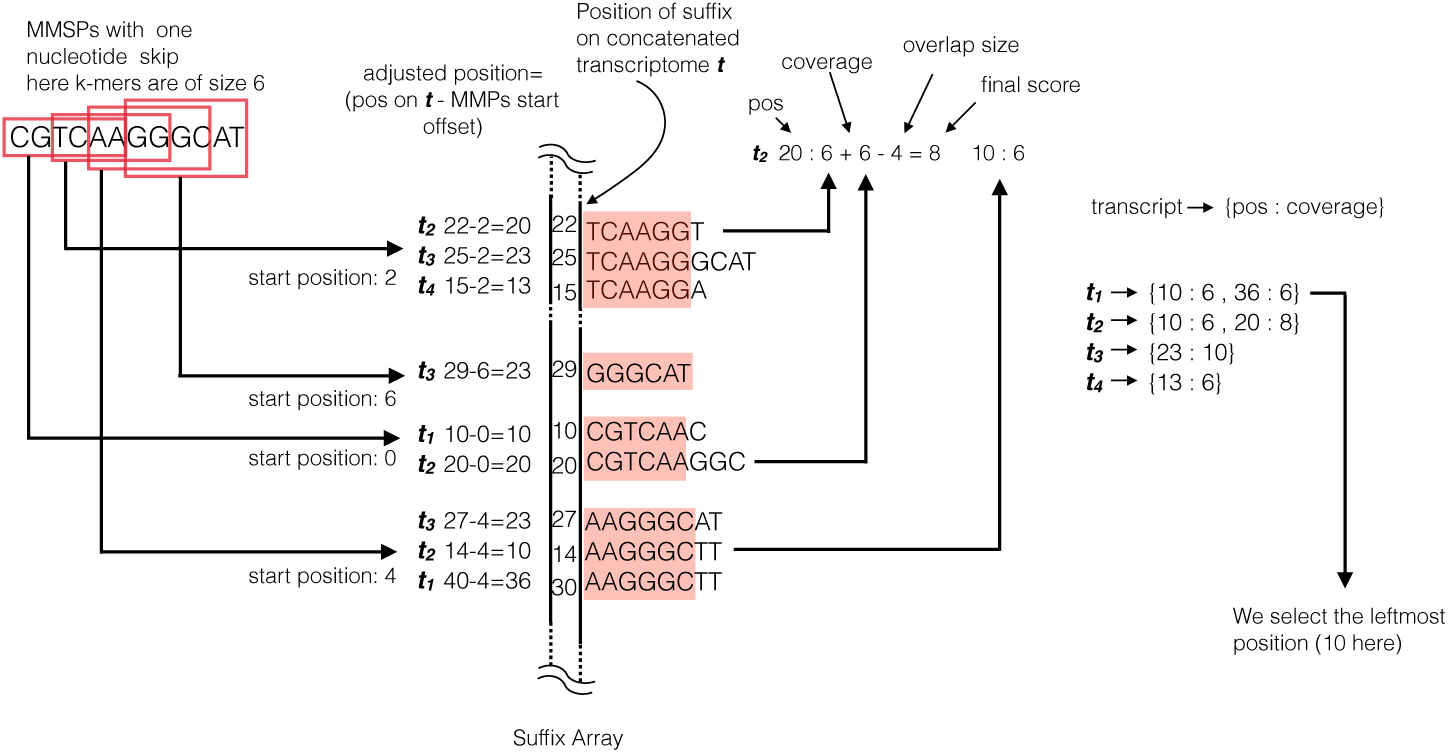
The MMSPs corresponding to a read, are derived from multiple suffix array intervals. Here, all MMSPs happen to be of length *k* as LCPs are of size *k*. The coverage scheme finds out the exact positions on each transcript by adjusting the starting position of the MMSPs. The total score takes into account the positions where matches overlap. The final position is chosen by selecting the locus with maximum coverage.

#### 2.3.2 Selecting the best candidate transcripts

Once the positional ambiguity within a transcript is resolved, the next step is selecting the best candidate transcripts from a set of mappings. Since mapping relies on finding exact matches, the length of the matched subsequence between the read and reference can sometimes be misguiding when comparing different candidate transcripts. That is, the transcripts with the longest exact matches do not always support optimal alignments for a read. At this point in our procedure, we follow the approach taken by many conventional aligners, and use an existing optimal alignment algorithm to compute the edit distance, by which we select the best candidate transcripts.

When performing alignment, we assume that a given read aligns starting at the position computed in the previous steps. This helps us to reduce the search space within the transcript where we must consider aligning the read, and thereby considerably reduces the cost of alignment. To align the read at a specific position on the transcript and calculate the edit distance between them, we use *M yer ′s* bounded edit distance bit-vector algorithm (Myers, 1999), as implemented in `edlib` (Šošić and Šikić, 2017). For a fixed maximum allowable edit distance, this algorithm is linear in the length of the read. We note that the bounded edit distance algorithm we employ will automatically terminate an alignment when the required edit distance bound is not achievable.

We remove all alignments with edit distance greater than a user-provided threshold. This is similar to the approach used by many existing aligners, and allows us to specify that even the best mapping for a given read may have too many edits to believe that it reasonably originated from a known transcript in the index. An appropriate threshold should be based on the expected error rate of the instrument generating the sequenced reads, and a very low threshold can, of course, lead to decreased mapping rate.

#### 2.3.3 Enhancement of quantification accuracy based on edit distance score

We investigated the effect of incorporating edit distance in downstream quantification. Since we integrated the selective-alignment scheme into the quantification tool *Salmon* (Patro *et al.*, 2017), the edit distance scores from selective-alignment can be used as a new parameter to *Salmon*’s inference algorithm.

In the framework of abundance estimation, we define the conditional probability of a fragment, *f*_j_, originating from a transcript, *t*_i_, as *P* (*f*_j_ |*t*_*i*_). Given the edit distance between the fragment and the transcript, we can incorporate this parameter into this conditional probability. *Soft* filtering introduces a new term in the conditional probability based on *d*_*i,j*_, which is the sum of the edit distances between the read ends of fragment, *f*_*j*_ and transcript, *t*_*i*_. We set this probability according to an exponential function, *P* (*a*_*j*_|*f*_*j*_ , *t*_*i*_) = *e*^*-*^4^*d*_*i,j*_^. The aggregate of threshold filtering and *soft* filtering can be described as follows:

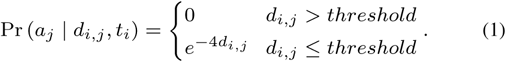

### 2.4 Shared LCPs prevents redundant alignments

Exploiting the common subsequences in the transcriptome is instrumental to the superior speed of fast mapping tools. Reads generated from exonic sequences common to multiple transcripts from the same gene or paralogous genes are the main source of ambiguous mapping. As we rely on the suffix array data structure to obtain the initial set of transcripts to which a read maps, there are cases where exactly identical reference sequences all act as mapping targets for the read. For a suffix array interval [*b, e*), we identify such common subsequences by examining the *longest common prefix* (LCP) of the interval. If the length of the LCP is equal or greater than the length of the read, then the actual alignment to the underlying reference at these positions will be identical.

Given the computationally intensive nature of alignment, this approach can be exploited to avoid the process altogether for some set of reference positions by simply reusing the alignment information from one read transcript pair and then passing it to other transcripts that share the LCP. As a proof of concept, we profiled the specific cases where such redundant alignments have been skipped in our algorithm. We observed (Table 1) that for almost half of the read-transcript pairs, the alignment process can be avoided. Note that if the read sequence shares a complete match with the common prefix, meaning that maximum mappable safe prefix length is equal to read length (i.e., the read matches the reference exactly at some set of positions), we can also bypass the Meyer’s edit distance algorithm call completely.

**Table 1.** The percentage of hits that skip the full alignment process on five different experimental samples, due to extension by the maximum mappable safe prefix (MMSP), or projection of duplicate alignments given the longest common prefix (LCP) sequences.

| Sample (SRR121) | 5996 | 5997 | 5998 | 5999 | 6000 |
| --- | --- | --- | --- | --- | --- |
| Skipped alignments | 50.36% | 54.85% | 47.92% | 48.06% | 50.80% |

## 3 Results

To evaluate the effectiveness of selective-alignment, we coupled it with the quantification tool *Salmon*. This enables us to measure the effect of different mapping/alignment algorithms on transcript-level quantification results directly, holding the statistical estimation procedure fixed. We also include *kallisto* in our benchmarks, which provides a perspective on pseudoalignment-based quantification. Furthermore, we compare the performance of selective-alignment with the recent, fast, abundance estimation tool *Hera* ^1^. We note, this is an early version of the *Hera* software (v 1.0), which is already performing well, but is subject to changes and improvements. We measure the Spearman correlation and Mean Absolute Relative Differences (MARD) of read counts as performance metrics when comparing the different methods.

### 3.1 Adversarial Synthetic Data

Genes with multiple isoforms are among the most challenging cases for aligning/mapping reads, since isoforms of the same gene share exonic sequences and are prone to a high degree of multi-mapping. Particularly complex regions of the transcriptome can pose a challenge to fast mapping algorithms, since many exact matches may occur at loci other than those which generate an optimal alignment. This can cause spurious mappings to mask true alignment locations, harming both sensitivity and specificity. Here, we generate an adversarial synthetic dataset which highlights potential mis-mapping problems. We restrict both the generation and assessment to multi-isoform genes. From the set of all multi-isoform genes in the human transcriptome (referred to as ground set), we selected a subset of transcript isoforms from which to generate reads. Through this mechanism, we ensure that only a fraction of the ground set of transcripts are truly expressed. Since the unexpressed transcripts share considerable sequence with the expressed transcripts, we expect a high rate of ambiguous multi-mapping.

The simulation procedure is randomized, and can be described as a two-step process. In the first step, we select a set of target transcripts (the foreground set) and quantify their abundances using reads from an experimental RNA-seq sample. In the second step, we generate synthetic reads from this set of estimated abundances and quantify the resulting data using the entire transcriptome.

To select the foreground set, we first examine each multi-isoform gene that produces protien coding transcripts, and select one such gene with probability *p*. Given the chosen gene, we select a candidate isoform with probability *q*. Following this protocol, we ensure that the number of truly expressed transcripts never exceeds 100 × *pq* percent of the number of transcripts in the ground set. For the two simulated datasets used here, 100 × *pq* is 30 and 60, respectively (*p* = 0.6, *q* = 0.5 and *p* = 1.0, *q* = 0.6). The motivation for this experimental set up comes from a previous analysis of the effect of expression “bleed through”^2^ on different quantification procedures.

To simulate data, *RSEM* (Li and Dewey, 2011) was run on sample `N12716_7` of the Geuvadis study (Lappalainen *et al.*, 2013), with the selected foreground set of transcripts (30% and 60% respectively) used as a reference to learn the model parameters and estimate true expression. The learned model is then used to generate 15 million, 75bp paired-end reads from each foreground set. These reads are then aligned/mapped to the set of all known transcripts from GRCh38.p10 using *Bowtie 2*, Hera, *kallisto*, *RapMap*, selective-alignment and STAR. Subsequently, transcripts are quantified by *Salmon* using the relevant alignments/mappings as input (except in the cases of *kallisto* and *Hera*, which implement their own quantification algorithm). The alignment mode of *Salmon* enables us to use *STAR* and *Bowtie 2* output as a direct input to the quantification module — thereby reducing variability due to differences in the underlying quantification model. To achieve the most sensitive alignment, *Bowtie 2* is run with the alignment options suggested for use with *RSEM* (Li and Dewey, 2011). For aligning reads to transcriptome using STAR we used the same option described in (Srivastava *et al.*, 2016). When processing alignments, *Salmon* was run with `--useRangeClusterEqClasses` (Zakeri *et al.*, 2017) and `--useErrorModel`. With selective-alignment, *Salmon* was run using `--useRangeClusterEqClasses`, `--softFilter` (discussed in Section 2.3.3) and an edit distance threshold of 4. *kallisto* was run with default parameters. Both the *Salmon* and *kallisto* indices were built with *k* = 25; *Hera* does not allow k-mer size as a user-defined parameter.

As displayed in Table 2, for the dataset where at most 30% of transcripts are truly expressed, the STAR, *Bowtie 2*, and selective-alignment-based methods perform better than the fast mapping approaches. Hera outperforms *kallisto* and quasi-mapping enabled *Salmon*, but does not perform as well as selective-alignment or the traditional alignment-based approaches. Presumably, this is because Hera uses a fast, banded alignment method to validate mappings. Thus, it benefits from similar improvements to precision as enjoyed by selective-alignment, though it doesn’t obtain the same improvements in sensitivity. Table 2 also demostrats that in terms of speed selective-alignment based salmon is comparable with ultrafast alignment-free approaches such as *kallisto* and quasi-mapping, which is considerably faster than alignment-based methods such as *Bowtie 2*.

**Table 2.** Performance of methods in terms of quantification accuracy on two foreground sets, 30% and 60%. quasi-mapping is the mapping approach used by RapMap. For STAR and Bowtie 2 we only record the timing for alignment step, and quantification is performed by Salmon. For kallisto and Hera timing includes the integrated quantificatios step alongside mapping.

| Method | Foreground | Spearman | MARD | time (s) |
| --- | --- | --- | --- | --- |
| STAR | 30% | 0.84 | 0.03 | 241 |
| Hera | 30% | 0.80 | 0.04 | 187 |
| kallisto | 30% | 0.79 | 0.05 | 54 |
| quasi-mapping | 30% | 0.78 | 0.05 | 52 |
| selective-alignment | 30% | 0.85 | 0.03 | 91 |
| Bowtie 2 | 30% | 0.86 | 0.03 | 1,158 |
| STAR | 60% | 0.84 | 0.06 | 244 |
| Hera | 60% | 0.83 | 0.06 | 119 |
| kallisto | 60% | 0.81 | 0.07 | 63 |
| quasi-mapping | 60% | 0.81 | 0.07 | 52 |
| selective-alignment | 60% | 0.85 | 0.06 | 85 |
| Bowtie 2 | 60% | 0.85 | 0.06 | 1,174 |

In the experiment where at most 60% of transcripts are truly expressed, the accuracy of all methods starts to converge, though the same trend of accuracy differences exists. Though we have designed these experiments to be adversarial in nature, they nonetheless raise an interesting point about how divergence between the true set of expressed transcripts and those considered during quantification might affect accuracy. Specifically, aligning/mapping against a larger and more comprehensive set of potential isoforms need not always yield superior results. When unexpressed isoforms share considerable sequence with those that are truly expressed, the probability of mis-assigning ambiguously mapping reads can increase. Though this is true regardless of how reads are aligned/mapped, alignment-based methods (and selective-alignment) seem less prone to mis-assignment in such cases.

### 3.2 Synthetic reads from human transcriptome

We have also explored the performance of different alignment-based and alignment-free methods and selective-alignment on the full human transcriptome. We follow the procedure described in (Bray *et al.*, 2016) to generate 30M, 75bp paired end reads using the *RSEM* simulator. Reads are mapped/aligned to the human transcriptome (Ensembl release 80 (Yates *et al.*, 2015)) with different methods, and then quantified by *Salmon*, *kallisto* or Hera. The Spearman correlation and MARD values for different methods are reported in Table 3. The performance of both alignment-based and alignment-free methods are similar to each other at the transcriptome-wide scale (and when not focusing on adversarial situations). *Bowtie 2*-based quantification seems to marginally outperform the mapping-based methods. Selective-alignment’s accuracy is very similar to that of *Bowtie 2*, but it requires considerably less time (it is similar to the fast mapping-based methods in this respect).

**Table 3.**
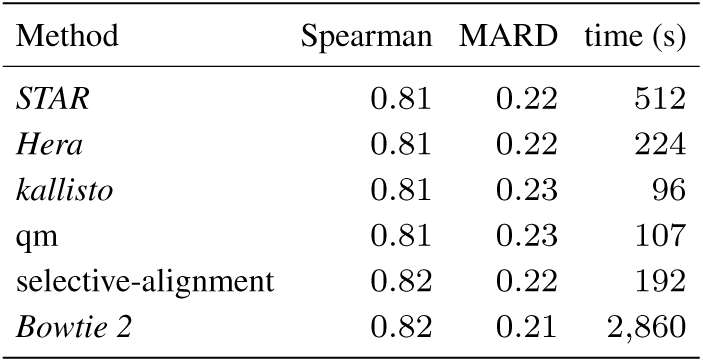
Quantification results with different methods for aligning/mapping reads on transcriptome wide synthetic data. The quantification is performed using same convention described in Table 2

Transcriptome-wide assessments on synthetic data, like that explored in this experiment, suggest that fast mapping-based methods generally perform well (and similar to alignment-based methods). However, small global differences in quantification accuracy at the transcriptome-wide scale tend to arise from larger differences in the quantification of particular transcripts (e.g., those where accurate mapping tends to be difficult, and where additional modeling fidelity is required to obtain accurate estimates (Zakeri *et al.*, 2017)). Such differences also arise, and tend to be somewhat larger, when analyzing experimentally-derived data, as we do in Section 3.3.

### 3.3 Experimental reads from human transcriptome

We have also benchmarked our proposed selective-alignment method, on experimental data from SEQC(MAQC-III) consortium (SEQC/MAQC-III Consortium and others, 2014) (NCBI GEO accession `SRR1215996`-`SRR1216000`). Each of five technical replicates consists of *~*11M, 100bp, paired-end reads, sequenced on an Illumina Hiseq 2000 platform. The options used for all methods are the same as those mentioned in Section 3.1. In Table 4, we compare the quantification results produced by different methods. Here, we note that we do not know the ground truth, and so we instead measure the overall concordance between different approaches. Each individual cell contains the average obtained across all five samples. High Spearman correlation and low MARD value between *Bowtie 2* and selective-alignment show that selective-alignment produces results more similar to *Bowtie 2* than to the other alignment-free methods.

**Table 4.**
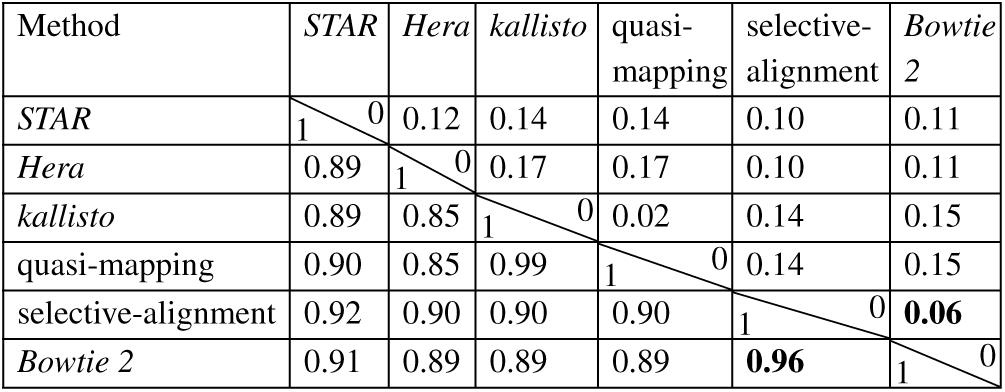
The Spearman correlation and MARDS between transcript abundances computed by all methods on experimental data. Each number is the mean on 5 different samples; the numbers in the lower left triangle of the matrix are the Spearman correlations and the ones in upper right are the MARD values.

## 4. Conclusion

Recently, fast mapping approaches have been developed for mapping RNA-seq reads to transcriptomes. Rather than generating full alignments, these approaches compute “mapping” information that is often sufficient f or a n umber o f g iven a nalysis t asks (e.g., transcript quantification (Bray *et al.*, 2016; Patro *et al.*, 2 017) o r metagenomic abundance estimation (Schaeffer *et al.*, 2017)). Yet, there exist scenarios where such mapping approaches can go awry; either failing, by the greedy nature of their procedures, to find t he t rue t arget o f o rigin o f a read, or by allowing spurious mappings to targets supported by exact matches that would nonetheless fail reasonable alignment scoring filters. Moreover, it is sometimes desirable to be able to produce, on demand, the edit distance or alignment that would result from a given mapping location. The recently-introduced *Hera* validates mapping quality using alignment, which resolves spurious mappings, though it still suffers a loss of sensitivity compared to traditional alignment. In this paper, we introduce a selective alignment algorithm that attempts to bridge the gap between these fast mapping algorithms and more traditional alignment algorithms. Selective-alignment improves upon both the sensitivity and specificity of these mapping algorithms while making very moderate concessions with respect to the computational budget. To achieve this level of efficiency, a number of algorithmic innovations were required, some of which may be of general interest. In the future, we hope to expand upon the notion of selective alignment even further, both by improving the algorithm and implementation, and by exploring use cases where selective alignment applies. Such situations are those where fast mapping approaches are inappropriate and traditional alignment approaches are too slow.

## Footnotes

1 https://github.com/bioturing/hera

2 https://cgatoxford.wordpress.com/2016/08/17/why-you-should-stop-using-featurecounts-htseq-or-cufflinks2-and-start-using-kallisto-salmon-or-sailfish/

